# Multivalent administration of dengue E dimers on liposomes elicits type-specific neutralizing responses without immune interference

**DOI:** 10.1101/2024.08.20.608851

**Authors:** Thanh T. N. Phan, Matthew G. Hvasta, Devina J. Thiono, Ruby P. Shah, Gisselle Prida Ajo, Wei-chiao Huang, Jonathan Lovell, Shaomin Tian, Aravinda M. de Silva, Brian Kuhlman

**Affiliations:** Department of Biochemistry and Biophysics, University of North Carolina at Chapel Hill, Chapel Hill, North Carolina, USA; Department of Microbiology and Immunology, University of North Carolina at Chapel Hill, Chapel Hill, North Carolina, USA; Department of Biomedical Engineering, University at Buffalo, Buffalo, New York, USA

## Abstract

The four serotypes of dengue virus (DENV1-4) are a major health concern putting 50% of the global population at risk of infection. Crucially, DENV vaccines must be tetravalent to provide protection against all four serotypes because immunity to only one serotype can enhance infections caused by heterologous serotypes. Uneven replication of live-attenuated viruses in tetravalent vaccines can lead to disease enhancement instead of protection. Subunit vaccines are a promising alternative as the vaccine components are not dependent on viral replication and antigen doses can be controlled to achieve a balanced response. Here, we show that a tetravalent subunit vaccine of dengue envelope (E) proteins computationally stabilized to form native-like dimers elicits type-specific neutralizing antibodies in mice against all four serotypes. The immune response was enhanced by displaying the E dimers on liposomes embedded with adjuvant, and no interference was detected between the four components.

## Introduction

Dengue viruses (DENVs), estimated to infect several hundred million people each year (1), continue to be a global health threat. The four DENV serotypes (DENV1-4) are genetically distinct but share similarities in structure, which generate both serotype-specific and cross-reactive antibodies (Abs) (2–7). Severe disease is most often observed following secondary infection with a DENV serotype that differs from the primary infection (8–11). In this scenario, there is evidence that cross-reactive Abs elicited during the primary infection fail to neutralize the secondary infection and mediate cellular uptake of the virus, enhancing the disease through a process termed antibody-dependent enhancement (ADE) (4, 12, 13). Traditional tetravalent vaccine approaches using live-attenuated viruses (LAV) have faced safety concerns due to uneven replication of the LAVs stimulating an unbalanced immune response to just one or two serotypes (14–18). The most advanced LAV DENV vaccine, Dengvaxia, is unbalanced and stimulates an immune response to mainly DENV4 (19). In this case, the vaccine mimicked a primary DENV4 infection for seronegative individuals. Consequently, young children without any dengue immunity who received this vaccine were at greater risk of hospitalization when exposed to DENV1, 2 or 3 infections compared to unvaccinated children (20, 21). Alternative vaccine approaches like subunit vaccines and virus-like particles, which are independent of virus replication, could potentially bypass this problem.

DENVs are enveloped viruses with a glycoprotein called the envelope (E) protein displayed on the viral surface. On the virion, the E protein forms 90 head-to-tail dimers which pack against each other to create a protein coat with icosahedral symmetry (22, 23). The E protein is the main target of the human antibody response (3). Neutralizing antibodies bind to a variety of epitopes on E protein, including quaternary epitopes which incorporate residues from both chains of the E homodimer (24–26). As subunit vaccines, wildtype soluble E proteins (sE) have multiple disadvantages: they are difficult to produce, are thermally unstable and do not form dimers at physiological conditions (27). Previously, we reported a set of amino acid mutations that stabilize soluble dimers across DENV1-4, increase expression levels, and improve their thermal stability (28, 29).

Cobalt-porphyrin phospholipid (CoPoP) liposomes are a vaccine platform that function as a nanoliposome vehicle for antigen display while enabling co-delivery of immunostimulatory adjuvants (30–32). CoPoP chelates cobalt ions within the lipid bilayer of liposomes, serving as an anchor to display proteins bearing polyhistidine tags (His-tags). His-tagged proteins rapidly and spontaneously associate with CPQ liposomes upon admixture in aqueous solution, with multiple copies of the protein presented on a single liposome. Besides CoPoP, also embedded in the lipid bilayer of the liposomes are the immunostimulatory vaccine adjuvants PHAD-3D6A, a form of synthetic monophosphoryl lipid A, and QS-21, a saponin, and together these liposomes are referred to as CPQ. These adjuvant components work synergistically to stimulate cytokine production and boost the antibody response. CPQ liposomes have been used in several vaccine studies (33–35). Most recently, a CoPoP vaccine displaying the SARS-CoV-2 receptor binding domain (EuCorVac-19) has successfully concluded phase II trial and progressed to a phase III trial (36–38).

In this study, we characterize the antibody response in mice of monovalent and multivalent formulations of stabilized E dimers and WT sE presented on CPQ liposomes. We also perform experiments with control liposomes that do not contain cobalt (termed LPQ) but include the same adjuvants as the CPQ liposomes, thereby isolating the impact of liposomal display. In this case, the E protein is not displayed on the liposomes and the liposomes just serve as an adjuvant. Our results show that the stabilized E dimers elicit more neutralizing antibodies than soluble WT E and that displaying the E dimer on CPQ liposomes improved both antibody quantity and neutralization for some DENV serotypes. From the multivalent vaccine studies, we found that the neutralizing response was type-specific with no interference between vaccine components. These results show that the stable E dimers are promising vaccine antigens and the tetravalent formulation with CPQ liposome is a potential candidate for a safe subunit vaccine.

## Results

### Stabilization of DENV1-4 E dimers

Soluble recombinant wildtype E proteins have low expression yields, do not form dimers at concentrations below 1 μM at 37°C, and do not bind tightly to neutralizing antibodies with epitopes that span across the E dimer (27, 29). To enable the use of the E protein as an effective subunit vaccine, we previously identified amino acid mutations that stabilize the E dimer and raise expression yields (28, 29). These E stabilized combinations (referred to as SC proteins) include mutations at the central αB interface (I2 or I9), the domain I-II hinge (S1, U6), and in the core of domain I (P4) (Fig. 1A, SFig. 1). Mass photometry experiments with SC variants of the four DENV serotypes show that all four proteins are primarily homodimers at a protein concentration of 50 nM, while WT DENV2 (Thai 16681), WT DENV3 (CH53489), and WT DENV4 sE (TVP-376) are almost exclusively monomeric (Fig. 1B,C). Purification yields were too low for the WT DENV1 E protein (WestPac74) to allow mass photometry experiments.

**Figure 1.**
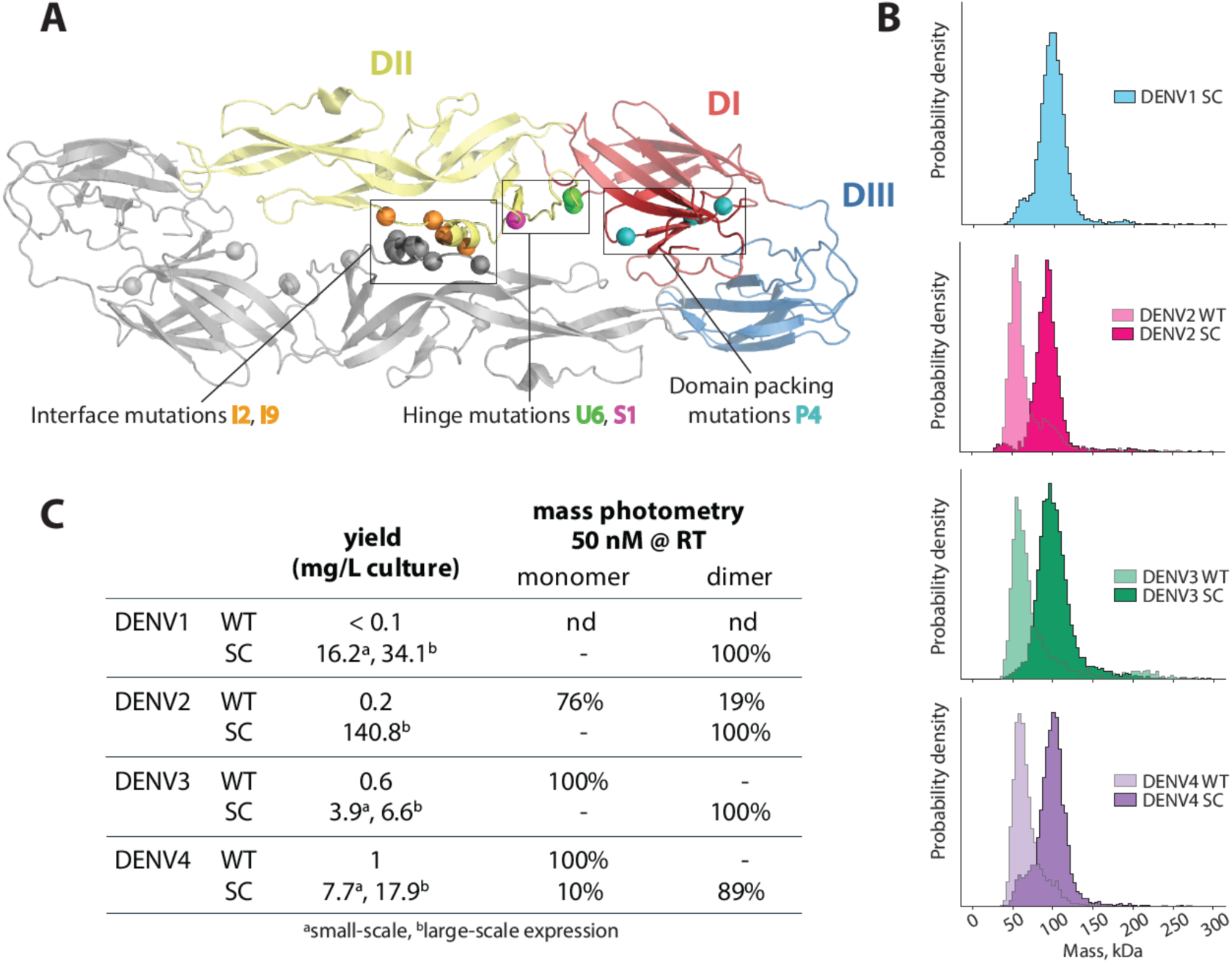
Biochemical characteristics of DENV soluble E with stabilizing mutations (sE SC). (A) Cartoon depiction of recombinant E homodimers with monomers (gray, colored) in head-to-tail orientation. Mutation sets at the dimer interface (I2, I9), DI/II hinge (U6, S1) and DI core (PM4) increased protein production and homodimer affinity. These mutations were applied to DENV1-4 sE to obtain stable dimers for all four serotypes. (B) Mass photometry histograms of DV sE WT and SC at 50 nM and room temperature. (C) Summary of yield and dimer stability of DENV sE. Yields and mass photometry data for DENV sE WT were obtained from (29), yields for DENV sE SC are from small scale and large-scale expressions in Expi293F; nd = no data, as we were unable to produce enough DENV1 sE WT for analysis.

### sE proteins are easily coupled to CPQ liposomes

For immunogenicity studies, we formulated sE proteins with liposomes to investigate the effects of antigen display and adjuvant on antibody response. The liposomes (LPQ and CPQ) are bilayer lipid vesicles containing the adjuvants PHAD-3D6A and QS-21 (Fig. 2A). In addition, both CPQ and LPQ contain CoPoP or its non-metal analog of porphyrin-phospholipid (PoP) respectively (SFig. 2A). CPQ liposomes act as an adjuvant and a display platform, whereas LPQ liposomes serve strictly as an adjuvant.

**Figure 2.**
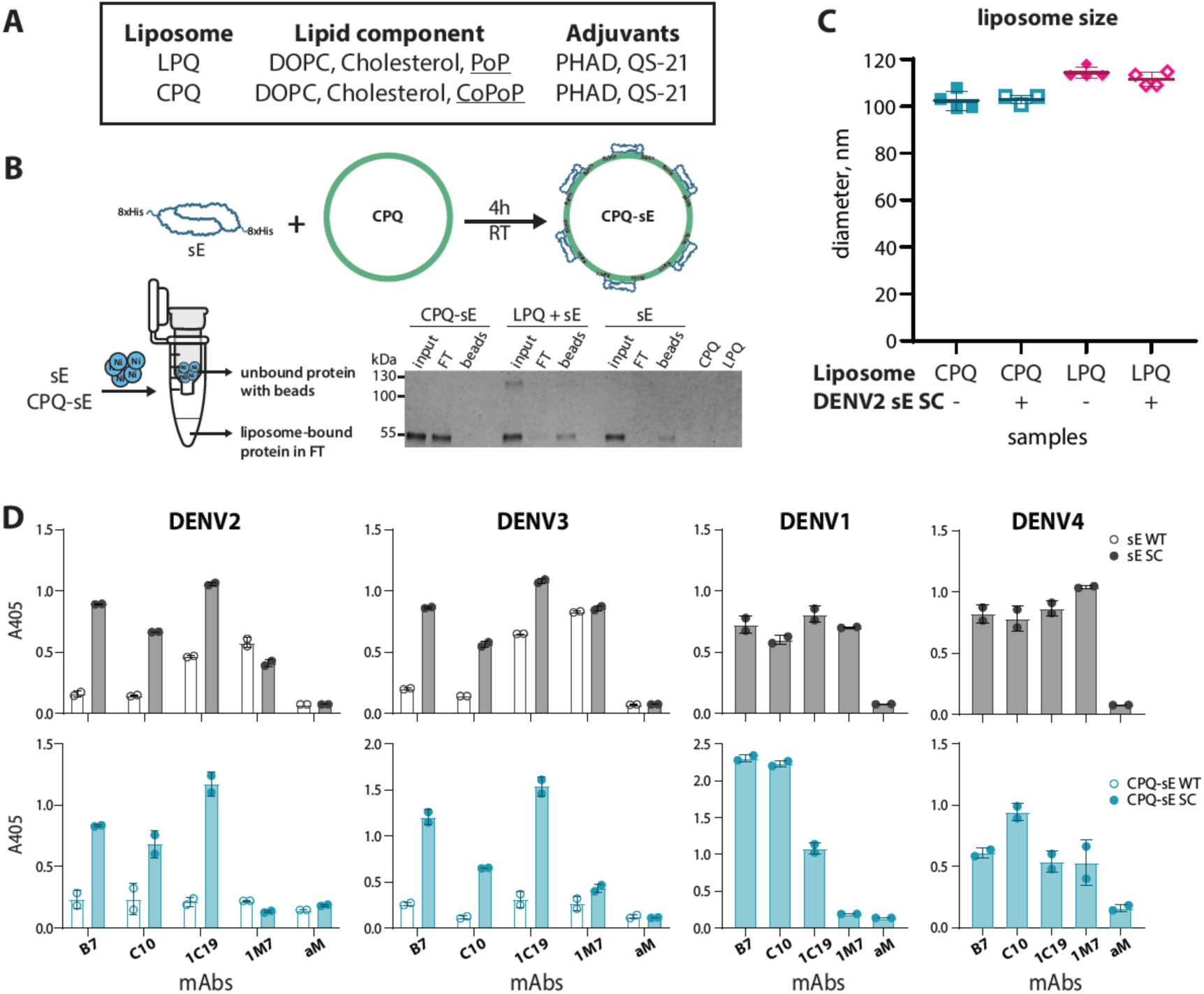
Characterization of sE-coupled liposomes. (A) Components in LPQ and CPQ liposomes. (B) Example conjugation scheme for sE SC and CPQ. sE proteins were mixed with CPQ at 1 sE : 4 CoPoP mass ratio for 4h at RT to conjugate antigens to liposomes. Complete conjugation results in 5-6 sE dimers per vesicle on average. Nickel (Ni) binding was used to assess conjugation efficiency. Free proteins (sE, ∼50 kDa) and CPQ-conjugated proteins (CPQ-sE) were incubated with Ni resin. Beads and flow-through (FT) fractions were analyzed by SDS-PAGE to determine bound and unbound protein. (C) Particle size characterization by dynamic light scattering (n=3 or n=4, mean ± sd). (D) Comparison between binding of DV sE SC (grey, solid) and sE WT proteins (grey, hollow) versus CPQ-coupled sE, CPQ-sE WT (blue, hollow) and CPQ-sE SC (blue, solid), to quaternary epitope Abs (B7, C10) and simple epitope Abs (1C19, 1M7). Anti-E Abs were conjugated with alkaline phosphatase (AP). AP-labelled anti-mouse IgG (aM) was used as negative control. Duplicate data represented as mean ± s.e.m.

sE proteins containing a C-terminal 8x His tag were incubated with CPQ liposomes for 4 hours at room temperature in the dark at a mass ratio of 1:4 sE:CoPoP (Fig. 2B). This mass ratio corresponds to 5-6 E dimers per liposome (SFig. 2B). As a control, proteins were also mixed with LPQ at the same ratio of sE:PoP. To assess the efficiency of coupling to LPQ and CPQ liposomes, we used nickel resin to pull out unbound proteins and separated them from liposome-bound sE via filtration (Fig. 2B, SFig. 2C). SDS-PAGE analysis of different fractions showed that sE proteins (∼50 kDa) mixed with LPQ samples were associated with the nickel beads, similar to soluble protein control, suggesting that there was no interaction between the His tag and LPQ. sE proteins incubated with CPQ liposomes were found in the flow-through fraction, indicating that the His tag of these proteins did not interact with nickel beads and that the proteins were fully bound to the CoPoP moiety.

The hydrodynamic diameter of the liposome samples did not change after protein incubation as evidenced by dynamic light scattering (DLS) (Fig. 2C). On average, CPQ liposomes (103 ± 4 nm preincubation, 103 ± 2 nm post incubation) remain slightly smaller than LPQ liposomes (114 ± 2 nm; 112 ± 3 nm). We observed no indication of protein-induced aggregation or fusion of lipid vesicles in any of the samples. From the DLS and nickel bead experiments, we concluded that sE proteins were completely attached to CPQ liposomes and that the integrity of liposomes was maintained after conjugation.

### Formulation with liposomes does not aBect Ab epitopes on sE proteins

We used well-characterized anti-dengue monoclonal antibodies (mAbs) to test if liposome-display interfered with sE structure and epitope accessibility (Fig. 2D). First, the binding profiles of unconjugated sE WT and SC at 37°C were obtained by coating soluble proteins on a nickel ELISA plate and probing with a panel of human mAbs. Consistent with the mass photometry data, DENV2 and DENV3 sE SC (grey bars) are stable dimers that exhibit stronger binding to E dimer dependent quaternary epitope binding mAbs B7 and C10 (24, 39) than WT proteins (white bars). The antibody 1C19 (40), which binds to the second domain of E (EDII), also bound more tightly to stabilized E dimers. In both serotypes, we found that fusion loop mAb 1M7 bound equally well to sE WT and SC. DENV1 and DENV4 sE SC also bound to both quaternary epitope mAbs and monomer epitope mAbs. These results show that the sE SC proteins are well folded and dimeric as they bind to both monomer and dimer epitope Abs, while the WT sE proteins only to monomer epitope Abs.

To assess whether coupling to CPQ liposomes affected epitope presentation, CPQ-sE samples were captured onto ELISA plates by human mAb 1M7. Similar to the unconjugated sE proteins, DENV2 and DENV3 CPQ-sE SC (Fig. 2D, blue shaded bars) show stronger binding to complex epitope Abs (B7 and C10) than CPQ-sE WT (blue hollow bars). As the binding signal of mAbs to CPQ-sE WT samples were low, we performed the same experiment using mouse mAb 4G2 as a capture antibody (SFig 3). Here, we observed that CPQ-sE WT had equivalent binding to 1M7 as the CPQ-sE SC samples, but weaker binding to dimer mAbs, mimicking the recombinant sE results. Given that the same amount of sE was fully coupled to CPQ liposome (SFig 2C), these findings suggest that liposome conjugation does not induce dimer formation of the sE WT protein and maintains dimer epitopes on sE SC. The binding of DENV1 and DENV4 SC to the same Abs were also not affected by CPQ coupling.

### Mice immunized with DENV CPQ-sE SC elicit antibodies that neutralize mature dengue virus

BALB/c mice were immunized intramuscularly with different formulations of proteins: DENV2 or DENV3 sE alone, a bivalent cocktail with DENV2 and DENV3 proteins and a tetravalent group with DENV1-4 sE (Fig. 3A). Within the monovalent and bivalent groups, we compared the immunogenicity of WT and SC proteins formulated with CPQ (displayed) or LPQ (non-displayed) liposomes. The tetravalent group consists of a cocktail of CPQ-sE SC from DENV1-4. In the multivalent experiments with CPQ liposomes, sE proteins from the various serotypes were coupled separately with liposomes and then combined for injection. With this approach, each individual liposome only displays E protein from a single serotype. We also included a bivalent mixture of the DENV2 and DENV3 stable dimers formulated with Alum as an adjuvant.

**Figure 3.**
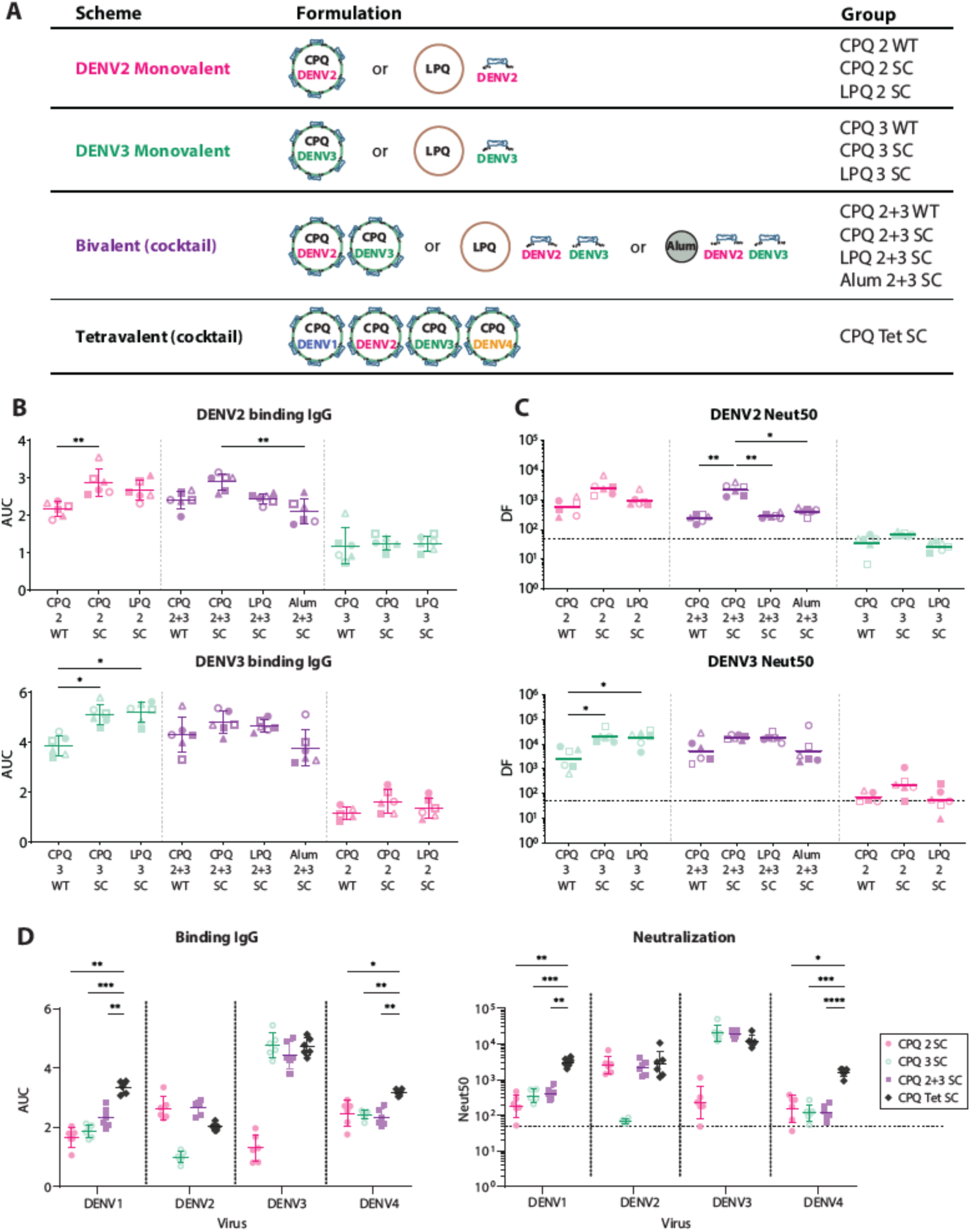
Virus-specific IgG titer and mature virus neutralization by mouse sera. (A) Mouse vaccination scheme: mice (n=6 per group) were immunized at weeks 0 and 4, with blood drawings on week 4 (pre-boost), 5, 8 and a final bleed at week 12. Groups include monovalent DENV2 or DENV3, bivalent DENV2 and DENV3 and tetravalent DENV1-4. (B) Placebo-subtracted total binding IgG titer to mature DENV2 (top) and DENV3 (bottom) represented as area under the curve (AUC) across groups, mean ± sd. (C) Mature virus neutralizing titers, DENV2 (top) and DENV3 (bottom), represented as the serum dilution factor at which 50% of the virus was neutralized (Neut50), mean ± sd. (D) Binding IgG (placebo-subtracted) and neutralizing titer against mature DENVs across monovalent CPQ-sE groups, bivalent groups and tetravalent CPQ-sE. The horizontal dotted line represents the limit of detection of the neutralization assays. Statistical analysis by a one-way ANOVA followed by a Tukey’s test (*p<0.05, **p<0.005, ***p<0.0005, ****p<0.00005).

All vaccinated mice generated Abs that bind the matched viruses. At week 12, mice immunized with DENV2 sE proteins produced similar amounts of binding antibodies to DENV2 in both the monovalent (pink) and bivalent formulations (purple) (Fig. 3B). Likewise, mice immunized with the DENV3 sE proteins produced similar amounts of binding antibodies to DENV3 in both the monovalent (green) and bivalent formulations (purple). Binding was largely type-specific, as the monovalent formulations did not generate large binding signals against the mismatched viruses. Within the monovalent groups, vaccination with the WT proteins induced lower levels of binding Abs than vaccination with SC proteins (p<0.05-0.005), while no significant binding differences were observed between the LPQ and CPQ formulations of the SC proteins.

For most animal groups, there was a strong correlation between the levels of binding IgG (Fig. 3B) and neutralization titers (Fig. 3C). Neutralization titers were measured with mature viruses produced in Vero cells with increased furin expression (Vero Furin cells). There was a robust type-specific neutralizing Ab response in the monovalent groups: i.e. all DENV2-vaccinated mice (pink) elicited high neutralizing Ab titer to DENV2 virus but not DENV3. Similarly, DENV3-vaccinated mouse sera only neutralized DENV3. Introducing 2 or more DENV serotypes in the formulations did not affect the strength of the response, evidenced clearly in the CPQ sE SC bivalent (purple) formulation.

Comparing across viruses, it was apparent that cross-reactive (CR) Abs alone could not generate robust responses (Fig. 3D). Only the tetravalent formulation (black) elicited strong binding and neutralizing titers against DENV1 and DENV4, outperforming both monovalent and bivalent formulations targeting DENV2 and DENV3. Strikingly, the tetravalent formulation induced approximately equivalent Ab levels as mono- and bivalent versions despite containing half the amount of each protein. Overall, the CPQ Tet SC formulation elicited a strong response against all four DENV serotypes with neut50 titers between 10^3^ and 10^4^.

In all cases, the sE SC formulations elicited stronger responses than those of sE WT. This may reflect more effective presentation of neutralizing epitopes, as evidenced by our binding studies with monoclonal antibodies (Fig 2). In comparing the LPQ and CPQ formulations, the effect of displaying antigens was most evident in the DENV2 neutralization assays. Both CPQ 2 SC and CPQ 2+3 SC groups elicited higher neutralizing titers than the corresponding LPQ groups. The same did not apply for DENV3-neutralizing response. LPQ and CPQ formulations were equally effective. Overall, the DENV3 neutralization titers were very high which may obscure differences between the LPQ and CPQ formulations. The bivalent CPQ SC formulation outperformed Alum (alhydrogel) adjuvant with the same protein antigens and doses.

### Neutralizing activity comes from type-specific (TS) Abs

To further probe the immune response to the various vaccine formulations, we performed depletion experiments to study the contribution of type-specific (TS) and cross-reactive (CR) Abs to virus neutralization (Fig. 4A). sE-coupled magnetic beads were used to remove all DENV specific Ab or just CR Abs from the mouse sera. For instance, with the bivalent CPQ 2+3 SC sera, we used DENV2 sE SC-coupled beads to pull out all DENV2-binding Abs, DENV3 sE SC-coupled beads to pull out all DENV3-binding Abs, and a combination of DENV1 and DENV4 sE to remove CR Abs that were elicited (Fig. 4B). After depletions, neutralization titers were measured with mature viruses produced in VeroFurin cells. The TS or CR nature of the neutralizing response is of interest because in humans TS Abs have been shown to be strongly protective and long-lasting, whereas CR Abs provide more breadth but have been implicated in weak protection and ADE. We used stable sE SC proteins to deplete CR Abs from WT-vaccinated mice as we were unable to produce all sE WT proteins.

**Figure 4.**
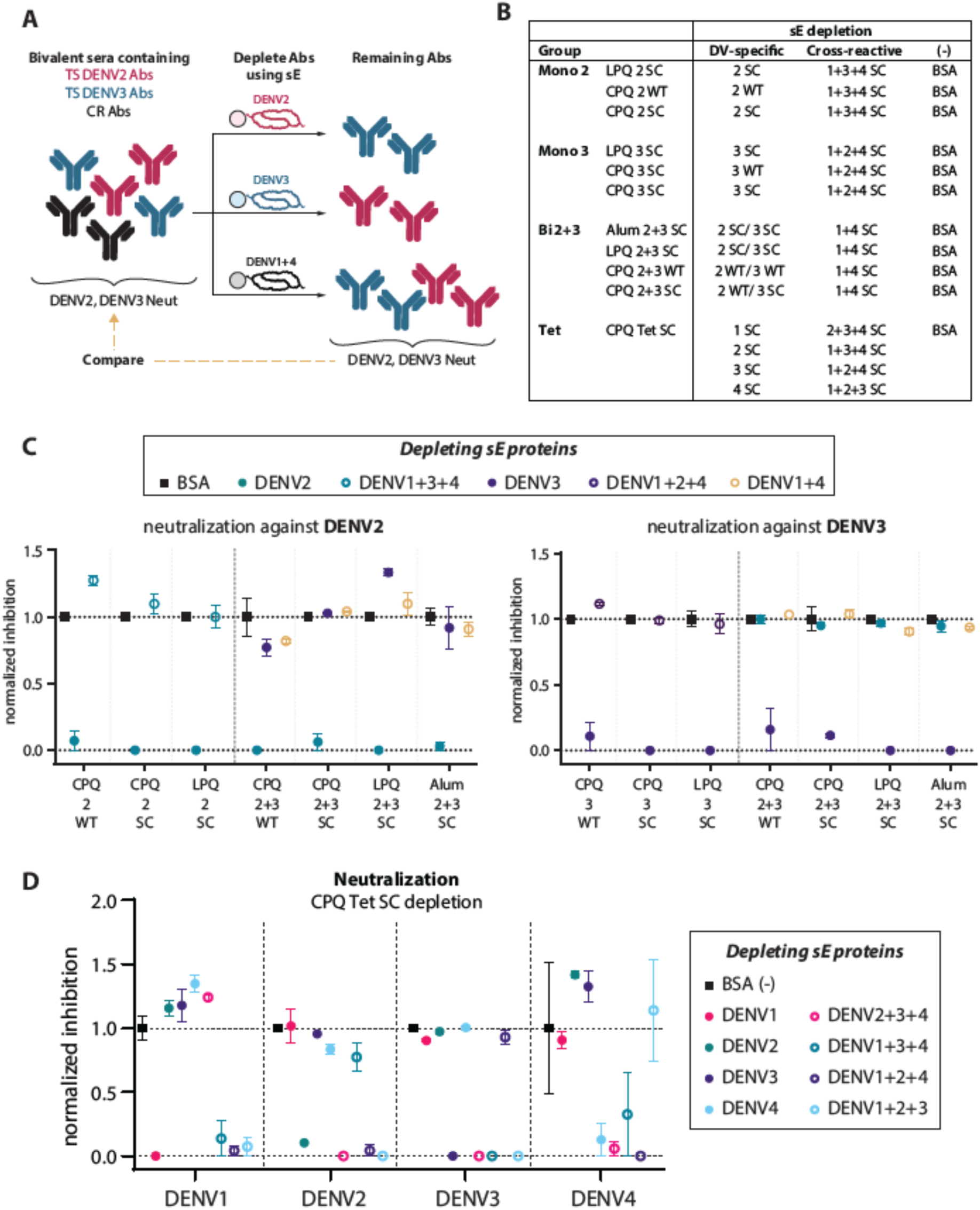
ENect of antibody depletion on DENV neutralization. (A) Representative schematic of a depletion experiment for mouse serum from bivalent DENV2+3 formulation. The serum is hypothesized to contain both type-specific (TS) Abs for DENV2 and DENV3, and cross-reactive (CR) Abs that can bind all four DENVs. Specific Ab components are pulled out of the serum by corresponding sE (DENV2, DENV3 or DENV1+4). The neutralizing response against DENV2 and DENV3 is re-evaluated after depletion to establish the protective role of TS or CR Abs. (B) Layout of the depletion experiment. Pooled serum from each group (monovalent DENV2, monovalent DENV3, bivalent DENV2+3 and tetravalent DENV1-4) was incubated with the corresponding vaccinating sE (WT or SC) to deplete all TS Abs and the remaining combination of proteins (SC only) to deplete CR Abs. BSA was used as a no depletion control. Each sample underwent 3 rounds of depletion. Neutralization of mature DENV2 & DENV3 by monovalent and bivalent pooled mouse sera (C) and DENV1-4 by tetravalent sera (D) at 1:420 dilution (DV4 at 1:250) after depletion by the vaccinating sE (solid circle) or the remaining sE (hollow circles). Neutralization was represented as virus inhibition of the depleted samples normalized to mean inhibition of the BSA control sample. Data represented as mean ± s.e.m. of duplicates.

For all vaccine formulations, the neutralizing response was maintained unless sE protein from the matched serotype was used for depletions (Fig. 4C, D). For monovalent DENV2 or DENV3 groups, removal of CR Abs by depleting sera with the other sE SC had little effect on neutralization (hollow teal or hollow purple circles). In bivalent formulations, we found that the mouse sera still maintained their DENV3 neutralization after the CR Abs were depleted either using DENV2 sE (solid teal circles) or DENV1 and 4 sE SC (yellow hollow circles). Similarly, there was little change in DENV2 neutralization when bivalent sera were depleted with DENV3 protein or DENV1+4 sE SC. Only when depleting with the matching sE to the virus (solid teal or purple circles), did the neutralizing response decrease significantly to background level. The same finding applies for the tetravalent formulation: the neutralizing response to each virus was completely diminished only if the matching serotype sE beads were present during the depletion. I.e., DENV1 neutralization was only abolished if DENV1 sE SC was used to deplete the serum and not DENV2, DENV3 or DENV4 individually or in combination. These results show that CR Abs generated by the mice play minor roles in the neutralization of mature DENVs and most of the neutralization activity comes from Abs specific to each DENV serotype.

## Discussion

The results of our study indicate that multivalent administration of DENV E dimers on CPQ liposomes in mice elicits TS neutralizing Abs against all four DENV serotypes with no interference. In most cases, the CPQ-sE SC formulations had higher neutralization titers than formulations based on WT proteins, LPQ adjuvant or Alum. The increase in immunogenicity was partly due to the antigen, which corroborates previous reports that sE dimers elicit higher IgG titers than WT and monomeric proteins (28, 41–43). The display of these proteins on the surface of liposomes may be stimulatory to B-cells, promoting cross-linking, expansion and maturation (44). The CPQ platform also offers a simple way to display different vaccine antigens. Overall, the boosted effect from multivalent display and protein engineering efforts to improve the E antigen open exciting areas for dengue vaccine research.

Previously, a subunit vaccine containing soluble recombinant wild type (WT) E proteins from DENV1-4 was evaluated (45–47). In a randomized phase I trial, this 3-dose tetravalent subunit vaccine (V180) produced DENV-specific neutralizing antibodies in flavivirus-naïve adults, but the tetravalent immunity decreased 14 months post dose 3, primarily due to a drop in DENV4 immunity to background level (46). In the preclinical phase of this study, mice vaccinated with 10 µg of each DENV E repeatedly elicited low amounts of neutralizing antibodies to DENV4 (47). Introducing these proteins in a tetravalent manner led to an overall decrease in DENV1-4 neutralization titers. Another study using DENV1-4 WT E adsorbed on PLGA nanoparticles also elicited anti-E neutralizing titer in mice, but the type-specific response was weaker for DENV1 and DENV2 (48). These studies highlighted the promise of E subunit vaccines; however, they also indicated that the antigens and delivery systems could be improved for a more balanced immune response.

From our mouse study, the tetravalent CPQ-sE SC formulation appears to be a promising subunit vaccine formulation. Compared to the previous subunit vaccines using WT E, CPQ Tet SC not only elicited a strong neutralizing response to all 4 DENV serotypes but also robust TS neutralization to each virus. It is likely that the neutralization response can be further balanced by adjusting the dosage of the proteins per injection or changing the ratio of proteins coupled to CPQ liposome. This could be a significant advantage compared to tetravalent DENV vaccines based on live attenuated viruses where it is challenging to balance viral replication and the immune response. Additionally, we performed binding and neutralization assays on fully mature DENVs produced in VeroFurin cells, which more closely resemble DENVs circulating in humans and are more difficult to neutralize than DENVs produced in Vero cells without overexpression of furin (49, 50). Most previous studies have performed neutralization assays with partially mature virus produced in Vero cells.

Further studies are needed to establish that the tetravalent CPQ-sE SC formulation will be an effective and safe vaccine in humans. Mice cannot naturally be infected with DENV, and thus are poorly suited to these types of studies. Studies in non-human primates (NHPs) can be used to evaluate the durability of the Ab response and perform challenge experiments with relevant strains of DENV. In the absence of human clinical data, Ab responses in NHPs have been a good predictor of vaccine outcome. A challenge study in NHPs using the tetravalent vaccine Dengvaxia (19) revealed that the amount of neutralizing Abs pre-challenge is negatively correlated to the magnitude of viremia, and that the vaccine only generates a robust TS response for DENV4, which has corroborated analysis of human sera post Dengvaxia administration (51). For the tetravalent CPQ-sE SC formulation to be a viable vaccine, it will be important to establish that the robust type-specific responses are reproducible in NHPs and humans.

In summary, we performed the first multivalent immunization studies with sE SC variants that were computationally stabilized to increase expression yield and dimer stability. Mutations at the interface, hinge region and domain I core of the E protein had stabilizing effects in all four serotypes. Displaying the proteins on the surface of liposomes did not perturb binding to monoclonal Abs that are known to neutralize the virus, including epitopes that span across the dimer interface. In mice, the liposome conjugated proteins elicited a strong neutralizing response that is specific to the serotype of sE on the surface. This study provides a foundation for further exploration of this platform as a tetravalent DENV subunit vaccine.

## Methods

### Cells and Viruses

Expi293 cells were used for protein expression. Monkey kidney epithelial Vero cells were used for virus neutralization assays. Soluble E protein sequences were based on DENV1 WestPac 74 (aa 1-394), DENV2 16681 (aa 1-394), DENV3 CH53489 (aa 1-392) and DENV4 TVP-376 (aa 1-394). Mature viruses produced in Vero cells overexpressing furin protease (VeroFurin cells) were used for binding and neutralization experiments (49, 50).

### Protein expression and purification

DNA plasmids were amplified from DH5α cultures and purified using endotoxin-free DNA extraction kits (Macherey-Nagel) as previously described (28). Purified plasmids were then transfected into Expi293 cells using Expifectamine reagent according to the manufacturer’s guidelines for small scale (25-50 mL) or large scale (200-500 mL) production. Cells were grown in a 37°C, 8% CO_2_ incubator at 130 rpm shaking. Media containing secreted proteins were harvested 64-72h post transfection, and clarified by centrifugation at 3,500 g (small scale expression) of 10,000 g (large scale expression) for 15 min. The supernatant was filtered through a 0.2 µm membrane and stored at 4°C until purification.

Purification of sE was done using Nickel affinity column followed by gel filtration. In brief, Nickel Penta resin (Marvelgent) was equilibrated with 10 column volume (CV) PBS and incubated with protein media in batch binding mode for 1 h or overnight at 4°C. Resin was washed with 6 CV (1M tris-HCl, 0.5 M NaCl, 25 mM imidazole, pH 8.0), 1 CV PBS + 25 mM imidazole pH 7.4 and 1 CV PBS + 50 mM pH 7.4. Protein was eluted with 1x PBS + 500 mM imidazole pH 7.4, concentrated and buffer exchanged into PBS + 10% glycerol pH 7.4 buffer for storage.

### Liposome synthesis

#### Materials

The following lipids were used to form CoPoP/PHAD/QS21 liposomes (CPQ); 1,2-dipalmitoyl-sn-glycero-3-phosphocholine (DOPC, Corden # LP-R4-070), cholesterol (PhytoChol, Wilshire Technologies), synthetic monophosphoryl Hexa-acyl Lipid A, 3-Deacyl (PHAD-3D6A, Avanti Cat # 699 855), and QS-21 (Desert King). Cobalt-porphyrin-phospholipid (CoPoP) or porphyrin-phospholipid (PoP).

#### Liposome preparation

CPQ liposomes were prepared by an ethanol injection method, followed by nitrogen-pressurized lipid extrusion in a phosphate-buffer saline (PBS) carried out at 55°C. Lipids were dissolved in 1 mL pre-heated 55°C ethanol for 10 min, then 4 mL of pre-heated PBS were added and incubated at 55°C for 10 min. The liposome extruder (Northern Lipids) was nitrogen pressurized and heated to 55°C with a pressure of near 200-300 PSI. Then the liposomes were extruded through 200,100 and 80 nm membrane filters stack for 10 times, followed by dialyzing in PBS at 4 °C twice to remove ethanol. Later, liposomes were passed through a 0.2 μm sterile filter, and QS-21 (1 mg/mL) were admixed with liposomes, the final concentration was adjusted to 320 µg/mL of CoPoP, 128 µg/mL of PHAD and 128 µg/mL of QS21 and stored at 4 °C.

### LPQ and CPQ liposome formulation

sE proteins were conjugated on CPQ liposomes following previously described protocols (33, 34). Specifically, the proteins were incubated with CPQ at a mass ratio 1:4 sE:CoPoP for 4h at room temperature in the dark. LPQ liposomes were also incubated with proteins in the same manner using 1:4 sE:PoP mass ratio. After the incubation, the samples were protected from light and stored at 4C.

### CPQ-sE conjugation eBiciency assessment by Ni binding and SDS-PAGE

25 µL sE or liposomes incubated with sE were incubated with either 5 µL 50% Ni resin slurry preequilibrated in PBS or PBS for 30 min at room temperature in the dark. The final protein concentrations were 40 µg/mL. The samples were mixed gently by pipetting every 10 min, and then transferred to a spin column and centrifuged for 1 min at 2,000 g to separate the Ni beads and the solution. Ni beads were resuspended using 27.5 µL PBS into a new tube. Flowthrough fractions and Ni beads were analyzed by reducing SDS-PAGE to assess for the presence of unconjugated sE in solution.

### Size analysis of LPQ and CPQ liposomes pre- and post-sE conjugation by dynamic light scattering (DLS)

DLS measurements were performed on a DynaPro Plate Reader II instrument in isothermal mode. Liposome samples were diluted in PBS to a final concentration of 100 µg/mL (Co)PoP. 2 mg/mL BSA in PBS was used as a control. The average hydrodynamic radius and polydispersity from 5 acquisitions (5s each) was plotted.

### Antibody binding to DENV1-4 sE and CPQ-sE by enzyme-linked immunosorbent assay (ELISA)

Binding studies were done according to previously published protocols (28). Soluble E proteins with a C-terminal His-tag (50 µL at 2 ng/µL) were directly coated on Nickel plates (Pierce, cat# 15142) in TBS buffer for 1h at 37°C. Plates were washed three times with 0.2% TBS Tween-20 (200 µL/well). Human anti-E monoclonal Abs were added to the plates at 2 ng/µL in blocking buffer (1x TBS buffer pH 7.4 containing 0.05% v/v Tween 20 and 3% non-fat dry milk) and incubated for 1h at 37°C, 300 rpm (50 µL/well). Plates were washed then incubated with goat anti-human IgG conjugated with alkaline phosphatase (AP) for 45 min at 37°C. After a final wash, wells were developed with pNPP. Binding was correlated with absorbance at 405 nm. The assays were performed in duplicate.

To capture liposome formulated sE proteins (CPQ-sE), human anti-E antibody 1M7 was coated in a 96-well plate (Greiner, cat# 655061) overnight at 4°C at 100 ng/well in 0.1 M NaHCO_3_ pH 9.6. Wells were blocked 1h at 37°C. CPQ-sE samples were diluted to 2 ng/µL sE in blocking buffer and added to the plate for 1h at 37°C, 300 rpm (50 µL/well). AP-conjugated human anti-E mAbs (using Abcam kit, cat# ap102850) were probed and detected by pNPP as described above. Additional binding studies were done with mouse mAb 4G2 as the capture Ab. For 4G2-captured samples, we probed binding with unlabeled human mAbs and goat anti-human IgG-AP (0.5 ng/µL).

### Mouse immunogenicity studies

Female BALB/c mice were purchased from Jackson Laboratory and used at 9–10 weeks of age. Mice were immunized intramuscularly with 60 μL of the vaccine formulation in PBS; 30 μL injection in each hind leg. For the monovalent formulations of DENV2 and DENV3 vaccines, each mouse received 2 μg sE (WT or stable dimer) formulated with LPQ or CPQ liposomes (n = 6). Bivalent formulations mixed 2 μg sE of DENV2 or DENV3 with the liposomes, and then mixed the liposomes (n=6). For the tetravalent formulation,1 μg of each sE was mixed with CPQ liposomes, and then the liposomes were combined. Control groups included 2 μg DENV2 and 2 μg DENV3 sE+125 μg Alum (n = 6), CPQ or LPQ alone (n=3). All groups were immunized with the same antigen formulation and dose at day 0 and 28, and serum samples were collected at indicated time points.

### Ethics statement

All experiments involving mice were performed according to the animal use protocol approved by the University of North Carolina Animal Care and Use Committee. The animal care and use related to this work complied with federal regulations: the Public Health Service Policy on Humane Care and Use of Laboratory Animals, Animal Welfare Act, and followed the Guide for the Care and Use of Laboratory Animals.

### Mouse serum binding ELISA to whole virus

The protocol is similar to the capture ELISA described above. Human 1M7 antibody was immobilized on Greiner plate (cat # 655061) in 0.1 M NaHCO_3_ buffer pH 9.6 (100 ng/well) overnight at 4°C. All incubations on the following day were done at 37°C with gentle shaking. Plates were blocked the next day with blocking buffer for 1h. Mature viruses cultured in Vero Furin cells were captured on the plate for 1h. Following washes, mouse serum was serially diluted in blocking buffer and added to the virus-captured plate (50 µL/well) and incubated for 1h. Plates were washed, and anti-mouse IgG-AP (Sigma-Aldrich) was added to the wells (50 µL/well) for a 45-minute incubation. After a final wash, signal was developed using pNPP, monitoring for absorbance at 405 nm. Assays were performed in duplicates. The binding IgG titer was calculated as the area under the curve (AUC) of placebo-subtracted binding signal against serum dilutions.

### Virus neutralization (FRNT) by mouse sera

Vero cells were seeded the night before at a density of 20,000 cells/well and should be at 80-90% confluency on the day of the experiment. Vaccinated mouse serum was serially diluted in growth medium (1% Antibiotic-Antimycotic (Gibco), 1% MEM Non-Essential Amino Acids Solution (Gibco), 1% L-Glutamine (Corning) in DMEM F12 (Gibco) containing 2% heat-inactivated FBS and incubated with virus for 1h at 37°C. The virus-serum mixture was then added onto cells (30 µL/well). After an hour incubation at 37°C, the mixture was removed from cells and overlay media (OptiMEM (Gibco) + 2% HI FBS + 1% Anti-anti + 1% (w/v) carboxymethylcellulose (Sigma-Aldrich)) was added to the plate (180 µL/well). Foci were allowed to develop for a varying amount of time. The incubation time for mature DENV1 is 48 h, DENV2 is 52 h, DENV3 is 52 h and DENV4 is 44 h. After the infection incubation window, overlay media was flicked off from the wells. Cells were washed with PBS and fixed with 4% PFA for 30 minutes at room temperature, and subsequently washed with PBS.

To visualize foci on fixed cells, cells were permeabilized with 1x PERM ((10X Perm buffer was diluted in MilliQ water. 10X Perm: 1% of bovine serum albumin (BSA, Sigma)) for 10 minutes at room temperature (50 µL/well) or overnight (100 µL/well). 1 ng/µL 1M7 antibody in PERM + 5% milk was added to each well for 1h at 37°C. Cells were washed with 1x PBS and incubated with 1:4000 anti-human IgG-HRP (Southern Biotech) for 1h at 37°C. After washing with PBS, 30 µL KPL True blue peroxidase substrate (SeraCare) was added to each well. After 15-30 min development at room temperature, wells were rinsed under a gentle stream of water and let dry. Wells were imaged using Immunospot camera and foci were counted using Viridot software (52). Focus reduction was calculated as: % neutralization = (foci_placebo_ – foci_serum_)/foci_placebo_*100%. Data plotted as average of duplicates and fitted to a sigmoidal dose-response curve.

### Depletion of sE-specific Ab from pooled polyclonal sera

sE proteins coupled to HisPur magnetic beads (ThermoFisher, cat# 88832) were used to pull out corresponding binding Abs. Beads were washed and equilibrated in PBS 0.05% Tween-20 containing 20 mM imidazole. Proteins were incubated with beads at 0.08:1 mass ratio in the same buffer. The final coupled beads were washed and stored in PBS pH 7.4.

To deplete anti-sE Abs from polyclonal serum, we used 32 µg proteins coupled to beads (400 µg beads) for each round of depletion. The beads were added to 96-well plate and washed with 100 µL PBS. A magnetic plate was used to pellet the beads and the plate was flicked to remove the solution. 170 µL 10-fold diluted pooled sera in PBS + 5% BSA was added to the corresponding depletion set up and beads were resuspended gently through pipetting. Any empty wells were filled with 150 µL PBS to minimize sample loss due to evaporation and the plate was carefully sealed. Sera and proteins were incubated for 1h at 37°C at 650-800 rpm in a thermomixer (Eppendorf) to ensure proper agitation. Once completed, the plate was spun at 4,000 rpm for 5 min and a magnetic plate was used to separate the beads from the sera. Sera was carefully aspirated from the wells to avoid mixing with beads. A negative control for depletion was performed by coating beads with BSA at the same mass ratio mentioned above. Each serum sample was depleted 3 times with the antigens listed in Fig. 4B. Depletions were validated using Ni ELISA checking for binding of polyclonal serum with the depleting antigen (e.g. if serum was depleted with DENV2 sE SC, depletion is complete if there is no longer binding of serum to DENV2 sE SC in Ni ELISA, SFig 4).

## Supplementary information

**SFig 1.**
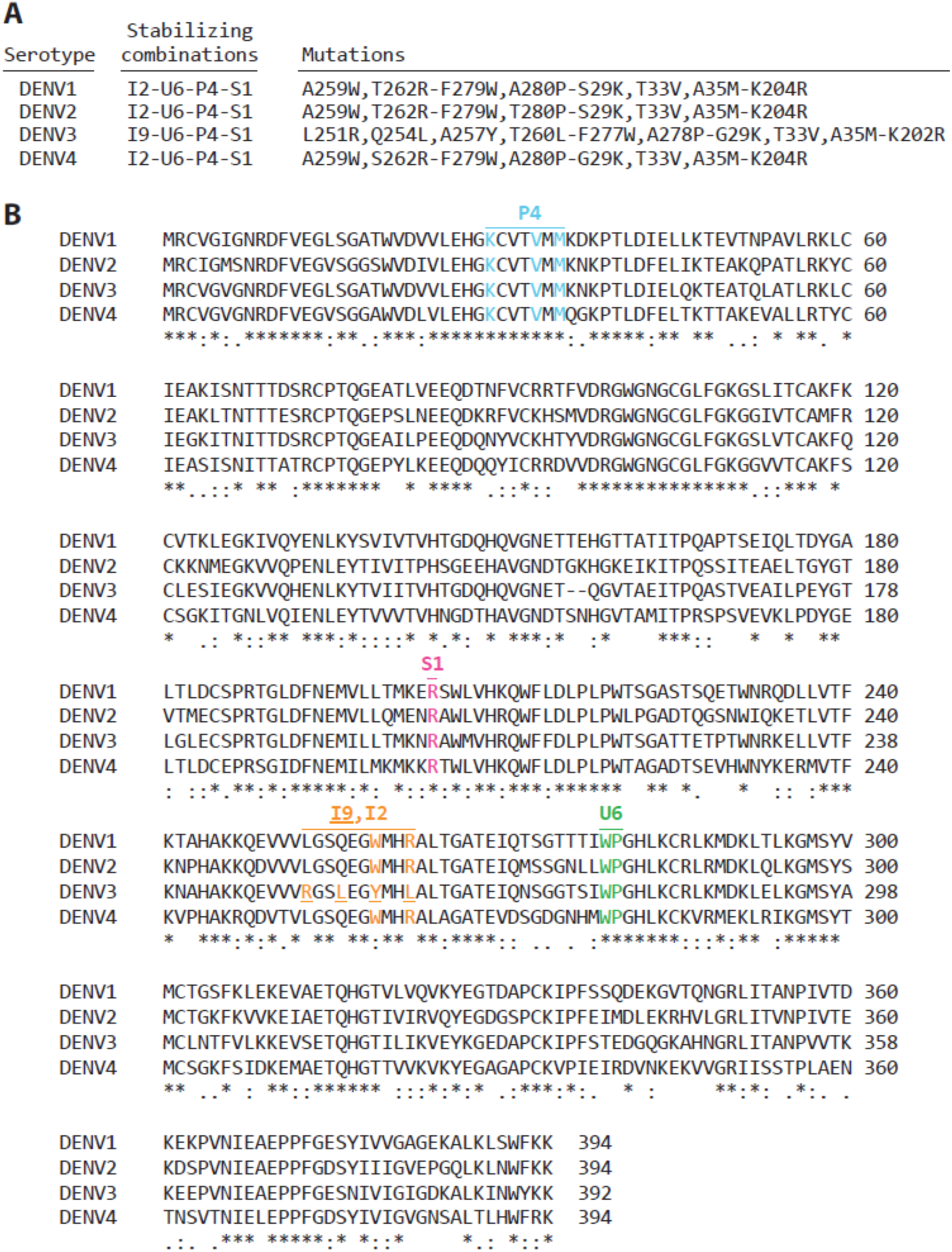
Sequences of DENV1-4 sE SC. (A) Mutations made in each serotype. (B) Sequence alignment of DENV1-4 sE SC. Amino acid sequences are from DENV1 WestPac 74, DENV2 16681, DENV3 CH53489 and DENV4 TVP-376. Stabilizing mutations are highlighted: PM4 in EDI (cyan), I2 and I9 at the central helix (orange), S1 (pink) and U6 (green) at EDI/II hinge.

**SFig 2.**
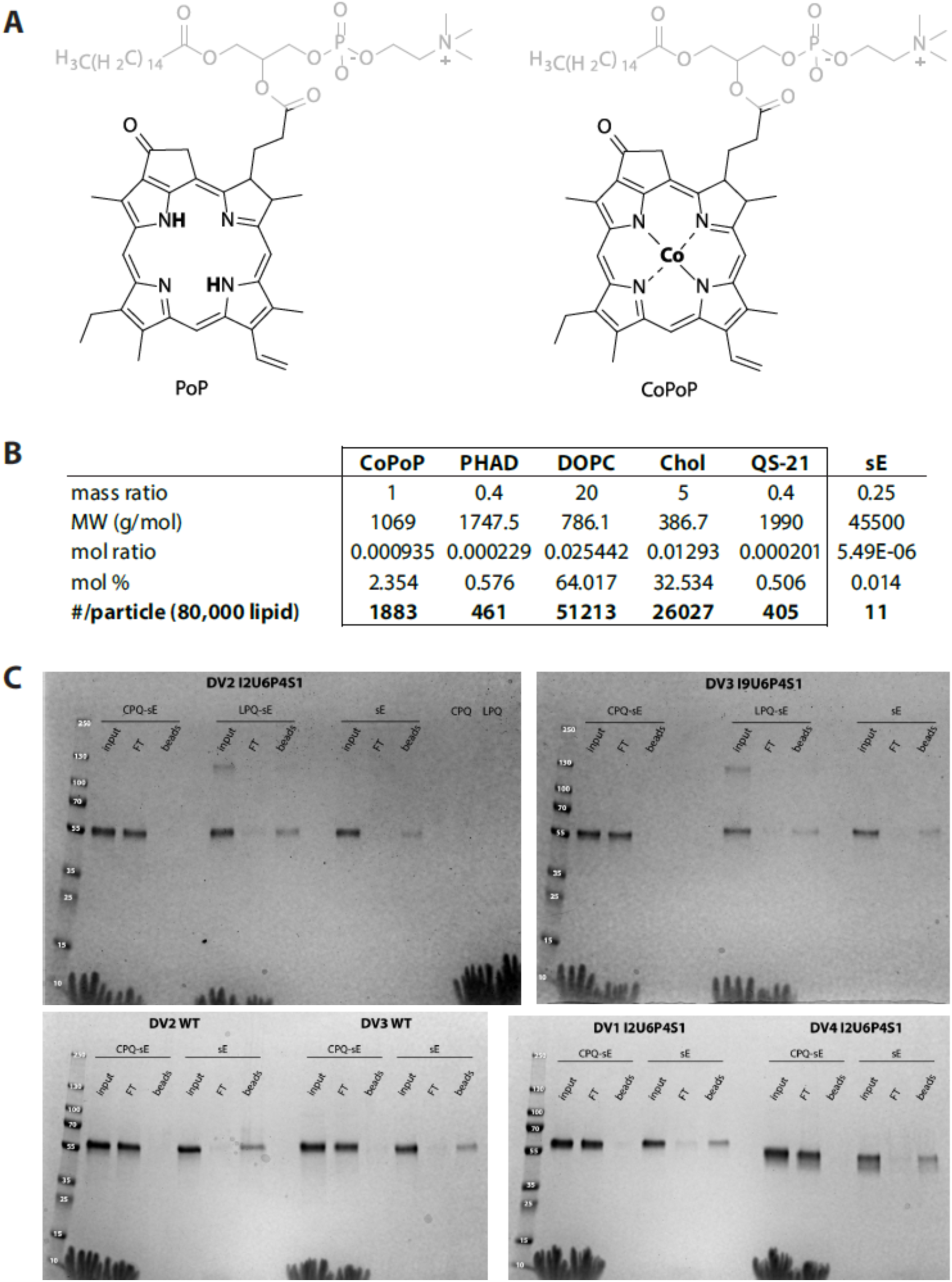
Coupling of sE to CPQ liposomes. (A) Chemical structure of PoP and CoPoP lipid moiety. (B) Stoichiometry calculation based on the coupling mass ratio. (C) SDS-PAGE analysis of unbound sE post conjugation to CPQ.

**SFig 3.**
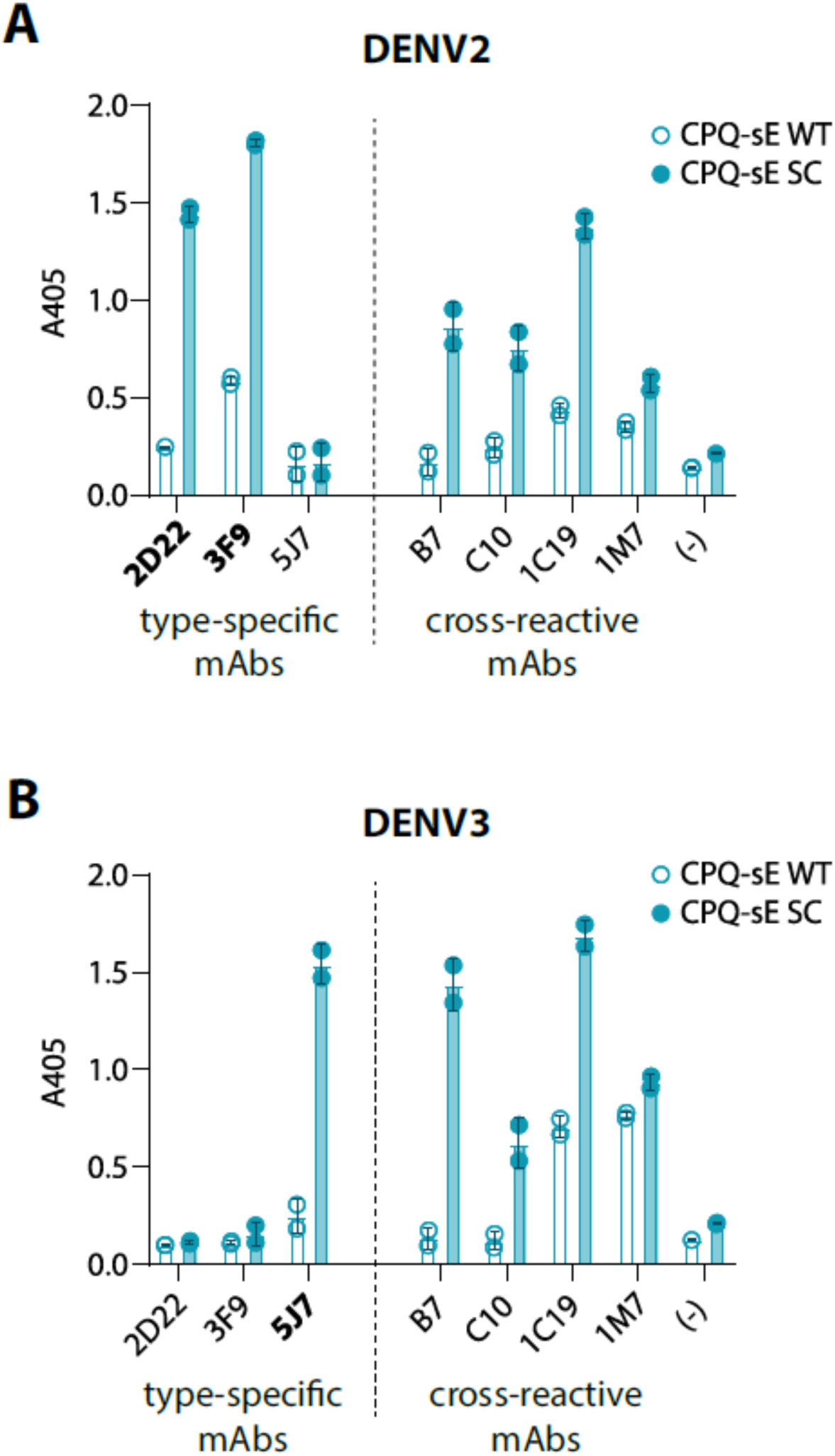
Binding ELISA of DENV2 and DENV3 CPQ-sE against a panel of type-specific and cross-reactive Abs. DENV2 (A) and DENV3 (B) CPQ-sE were captured by mouse mAb 4G2 and tested against human anti-E mAbs. Primary Abs were detected using AP-labelled anti-human IgG. (-) samples contained no primary Ab. Type-specific Abs for each serotype are bolded.

**SFig 4.**
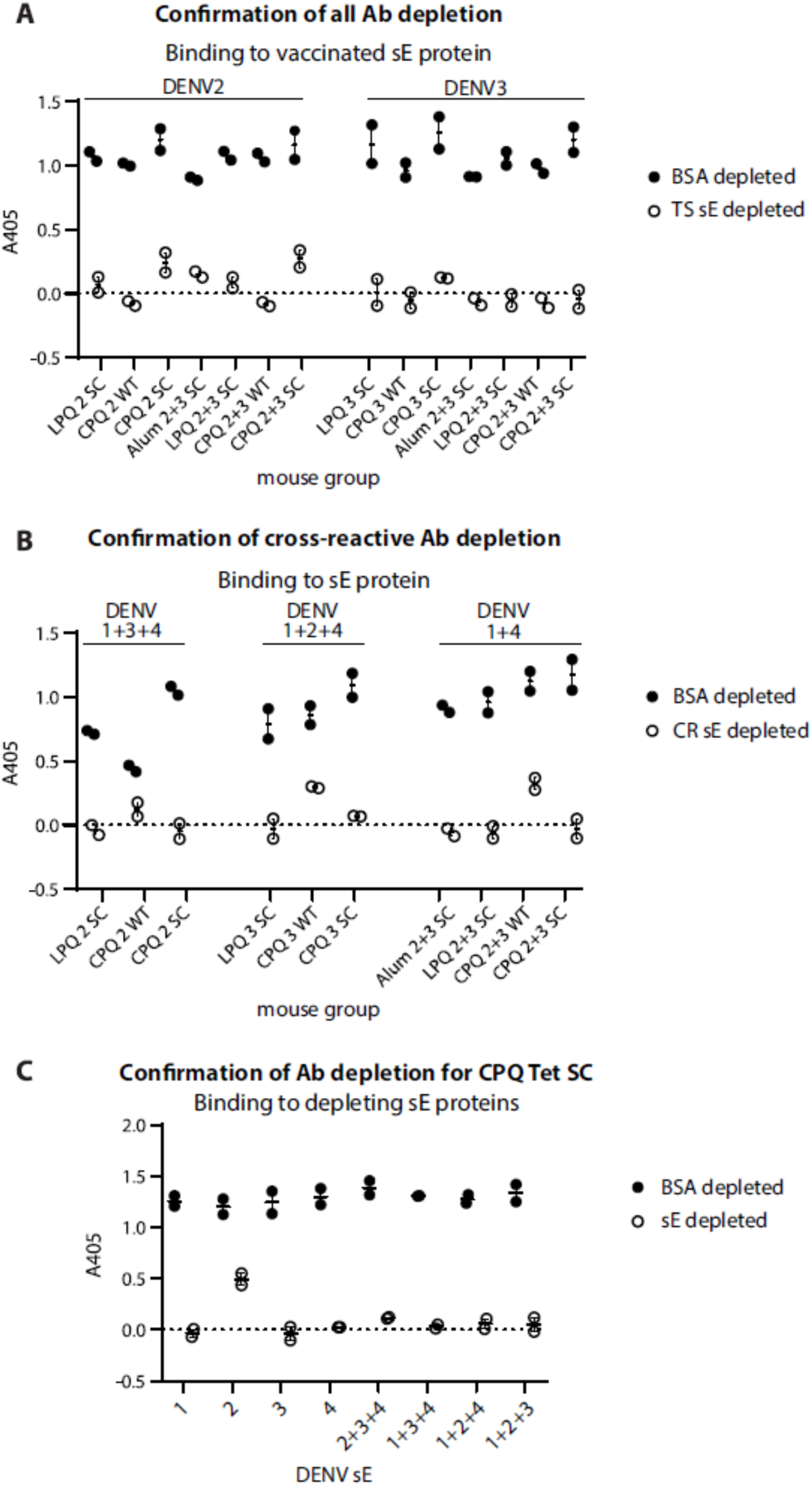
Confirmation of sE depletion by ELISA. Sera was pooled in each group for sE depletion and was incubated with sE-coupled or BSA-coupled beads to remove different Ab populations. Depletion was confirmed completed at 1:200 serum dilution by Ni ELISA to the matching depleting proteins for type-specific Ab depletion (A) and cross-reactive Ab depletion (B). (C) Type-specific and cross-reactive Ab depletion confirmation for CPQ Tet SC group. The dashed line represents no serum control

## Acknowledgements

We would like to thank John Forsberg at the UNC Protein expression & purification core for providing WT proteins.

## Author contributions

T.T.N.P, A.d.S and B.K. conceptualized the work. S.T., M.H. and G.P.A. performed animal studies. T.T.N.P., M.H. and G.P.A biochemically characterization of proteins. W.H. and J.L. synthesized liposomes. T.T.N.P., D.T., and R.S. contributed to serological analysis. T.T.N.P. created the first draft of the manuscript.

## Materials & correspondence

Any requests for materials should be addressed to T.T.N.P. and B.K.

